# A mathematical kinetic model of memory in *Bacillus subtilis* spore germination

**DOI:** 10.1101/2025.10.14.682329

**Authors:** Chris G. de Koster, Xiuping Lin, Yong-qing Li, Stanley Brul, Leo J. de Koning, Peter Setlow

## Abstract

Dormant *Bacillus subtilis* spores germinate through interaction of germinants with germinant receptors (GRs). Subsequently, the activated GR signal is transduced to SpoVA proteins that constitute transport channels. The opening of the SpoVA channel then leads to calcium dipicolinic acid (CaDPA) release and then completion of spore germination. Spores are known to exhibit memory in germination, as spores given an initial short germinant pulse respond more readily to a second pulse. We have developed a mathematical model to identify a minimal reconstructed molecular network that is crucial for germination kinetics leading to memory of germinant exposure; the model reproduces experimental double germinant pulse germination curves. Analysis of the reconstructed network indicates that a minimal set of inactive and active GRs and a SpoVA channel in three states - closed inactive, closed active and open - is needed to reproduce memory. Spore germination memory is introduced in the network by the activation and deactivation rates of GRs, and by the interplay between activation of closed SpoVA channels and their rate of opening and closing.

**Author summary:** Dormant bacterial spores return to life in germination when exposed to germinants which are recognized by germinant receptors (GRs). Spores have memory of germinant exposure, such that more spores germinate after a 2^nd^ germinant pulse than after the 1^st^ one, if intervals between pulses are short. In this work, a mathematical model has been developed that reproduces experimental memory behavior. Computer simulations of the model reveal the crucial factors in spore memory of germinant exposure as: i) active and inactive GR states and ii) a membrane channel for a major spore core molecule, Ca-dipicolinic acid (CaDPA), with the channel existing in closed inactive, closed active and open states, all of which are indispensable for spore memory. The model will be part of an iterative cycle of modelling of network dynamics and generating hypotheses for further experimentation.

## Introduction

Spores of bacteria of *Bacillus subtilis* are formed in sporulation, and are dormant and with an overall structure quite different from that of a growing cell (1). From the outside in, spores contain an exosporium in some species (but not *B. subtilis* spores!), then a layer of protein termed the coat, then an outer membrane above the spore’s peptidoglycan cortex that contains muramic acid lactam, next a layer of germ cell peptidoglycan with no muramic acid lactam, then an inner membrane (IM) encircling the spore core that contains all nucleic acids, most spore enzymes and a high level of the 1:1 chelate of Ca^2+^ with pyridine-2,6-dicarboxlic acid (CaDPA, ∼ 10% of spores core dry wt), as well as low levels of core water, as low as 20% of core wet weight (1). Spores are extremely resistant to all manner of treatments, have no ATP, but can survive harsh environmental conditions for many years (1–3). However, when nutrients become available and conditions are favorable for growth, spores can germinate and outgrow into vegetative cells via a sequence of molecular processes and signaling through pathways and networks (4–7). In the 1^st^ stage of germination the spore probes its environment for the availability of nutrient germinants. Upon interaction of germinants with germinant receptors (GRs) in the IM, spores germinate. *Bacillus* GRs generally have 3 subunits, A, B and C and these are gated ion channels with other ion channels participating (8–12). The binding of germinants, in particular to GR’s B subunits, leads to release of the spore’s huge depot of (CaDPA). This release then triggers the activation of two cortex-lytic enzymes (CLEs), CwlJ and SleB. In the 2^nd^ stage of germination, the CLEs hydrolyze the large spore cortex peptidoglycan (PG) layer, as these enzymes recognize muramic acid lactam which is only present in cortex PG (1–4). Cortex degradation then allows core expansion with full hydration of the spore core which allows resumption of normal metabolism, macromolecular synthesis, and finally outgrowth into a vegetative growing cell.

The GR protein assemblies GerA, GerB and GerK in *B. subtilis* are each encoded by homologous tricistronic operons *gerA*, *gerB* and *gerK* (1–4). GR subunits A and B are integral membrane proteins that are embedded in the spores’ IM through multiple predicted α-helices. Subunit C is a peripheral membrane protein, attached to the IM by a diacylglycerol anchor (13). The GerA GR responds to L-alanine or L-valine, the GerB and GerK GRs cooperatively respond to a mixture of L-asparagine, D-glucose, D-fructose and K^+^ (AGFK). A GerB variant, GerB*, germinates in response to L-asparagine alone. Average wild-type GR spore copy numbers have been determined by Western blotting and are ∼2,500 total GRs/spore, with GerA at ∼1,100 molecules/spore; and GerB and GerK each at ∼700 molecules/spore (14). As expected, GR levels in spores affect the germination rate, and GR overexpression increases rates of spore germination by these GRs (15). All *B subtilis* GRs co-localize in the spores’ IM in a cluster termed the germinosome (16), and germinosomes have also been found in other *Bacillus* species (17). The GerD protein co-localizes with GRs in the germinosome and is essential for germinosome formation, presumably acting as a scaffold, although the germinosome structure has not been determined. GerD is also essential for normal GR-dependent spore germination, as *gerD* spores germinate extremely poorly with L-valine via GerA or AFGK via GerB plus GerK (18). Like GR C subunits, GerD is also likely attached to the IM through an N-terminal diacylglycerol anchor attached to an N-terminal cysteine residue (18).

The signal generated upon GR activation, likely involving cations released from the spore core, is transduced to SpoVA proteins that make up a channel for CaDPA in the spore IM. However, the signaling mechanism is not fully understood, and no proteins that act in between the GRs and the SpoVA channel protein complex have been identified. The *Bacillus subtilis spoVA* operon encodes SpoVAA, SpoVAB, SpoVAC, SpoVAD, SpoVAEb, SpoVAEa and SpoVAF proteins. SpoVAA, SpoVAB, SpoVAC, SpoVAEb and SpoVAF are predicted to be integral membrane proteins based on their primary sequences, and SpoVAD and SpoVAEa are hydrophilic, soluble proteins that are on the outer IM surface (19,20). The copy numbers of SpoVAD and SpoVAEa have been determined by western blotting a ∼6,000 - 6,500 and 750 molecules/spore, respectively (19,20). SpoVAC alone functions as a mechanosensitive channel for small molecules such as CaDPA, with several treatments including membrane tension and the non-GR-dependent germinant dodecylamine opening this channel in vivo and in vitro (4,21,22). While the arrangement of SpoVA proteins in a supramolecular structure is not known, it seems most likely that the SpoVA proteins are subunits of a protein assembly in the spore’s IM that constitutes a channel for CaDPA movement into the developing spore during sporulation and out of the germinating spore (23). In contrast to GRs which are in the multiprotein germinosome, SpoVA proteins are more evenly distributed over the spore IM (16). Opening of the CaDPA channel is triggered not only by GR activation, but also by dodecylamine or hydrostatic pressures of ≥ 4000 atmospheres (4). Notably, while there is slow leakage of CaDPA from spores after germination is begun (24), almost all CaDPA is released in a very short time and in parallel with water uptake into the core (4).

In the 2^nd^ germination stage, CaDPA activates the CLEs which hydrolyze the cortex PG. CwlJ is activated first by CaDPA released from the spore core and SleB is activated only after substantial CaDPA release, perhaps by changes in cortex PG structure accompanying CaDPA release and water uptake. CwlJ can also be activated by exogenous CaDPA. The structure of the dormant spore cortex PG and physical effects due to elastic and contractile properties of the cortical PG sacculus presumably affect the susceptibility of opening of SpoVA channels, as does crosslinking in the spore coat which continues even after spores are released from sporangia (25,26). Cortex hydrolysis degrades the cortex PG macromolecular network restricting expansion of the spore core, thus enabling further uptake of water. As core water content increases, eventually full core hydration allows onset of metabolic reactions including biosynthesis of molecules needed for cell growth.

Spore germination is a heterogeneous process, as in populations some spores germinate rapidly and some extremely slowly (27–31). This heterogeneity is due primarily to a variable time period between the application of a germinant stimulus and the onset of rapid CaDPA release from the spore core. This time interval is denoted as the lag time *T*_lag_; all CaDPA is released by *T*_release_, and cortex PG hydrolysis and core swelling are complete by *T*_lysis_. Spores that are exposed to germinant molecules for a short time interval can be committed to germination at time *T*_c_ even when the germinants are removed and further binding to GRs is blocked (28–31); the slow leakage of 15-25% of CaDPA begins at or close to *T*_c_ and this slow release ends at *T*_lag_, when rapid CaDPA release begins (24). *T*_lag_ values are decreased by spore heat activation, higher GR levels and higher levels of GR-dependent germinants up to saturation (19). Spores with ∼4 fold higher SpoVA copy numbers have lower *T*_lag_ values in GR germination than those for wild type spores (15). The times needed for rapid release of all CaDPA remaining at *T*_lag_ (Δ*T*_release_ = *T*_release_-*T*_lag_) (0.5-3 min), and for cortex hydrolysis and core swelling (Δ*T*_lysis_ = *T*_lysis_ - *T*_release_) (10-20 min) are generally shorter than *T*_lag_. GR levels and nutrient germinant concentrations do not appreciably affect Δ*T*_release_ or ΔT_lysis_ values. However, loss of the CwlJ CLE, but not SleB, markedly decreases rates of CaDPA release during spore germination.

Notably, spores were found to exhibit memory of prior germinant exposures (30,32,33). Thus spores given a brief exposure to a germinant that activates the GerA or GerB* GRs responded more readily to a 2^nd^ short germinant pulse. However, the response to the 1^st^ germinant pulse diminished when the time interval in between pulses was increased (30, 32), indicating that there is memory in the germination system. Furthermore, nonnutrient germinants such as GR-independent CaDPA or dodecylamine, which do not require GRs, exhibit memory either alone or in combination with nutrient germinants (32), suggesting that the SpoVA channel proteins are the most likely candidates for memory storage while GRs are the candidates for memory generation. It has been proposed that spore memory is primarily stored in metastable states of SpoVA proteins (32), and/or that the spore’s electrochemical potential plays a role in enabling this memory (34). However, the involvement of electrochemical potential in the memory of germinant exposure appears unlikely as it was found that Thioflavin-T cannot report on electrochemical potential of germinating spores (33), but mechanisms underlying spore germination memory are not clear.

The pioneering work on mathematically modeling germination kinetics and heterogeneity was conducted by Woese et al. in 1968 (35). Spore germination was assumed to result from the accumulation of an unknown substance P, with its production rate being proportional to the number n of activated germination “enzymes”. By assuming that a threshold amount of P was required for initiating germination, it was predicted that time-to-germination of a spore with n active enzymes is proportional to 1/n. Furthermore, by assuming that the fraction of spores with n active enzymes in a population follows a Poisson distribution, this simple model explained spore heterogeneity and germination kinetics. Recently, Woese’s work was extended to a comprehensive mathematical model to describe heterogeneity, commitment, memory, and kinetic CaDPA release of spore germination (36), in which each spore in a population possesses m functional GRs following a statistical distribution, the binding of nutrient germinant activates a number of GRs and produces a time-dependent germination trigger signal. It was assumed that when the trigger signal is above a critical value, the spore is committed to germinate. This commitment results in the opening of the SpoVA channel, leading to the release of CaDPA and full spore germination (36). This model’s predictions fitted very well with the experimental data of heterogeneity, commitment, memory, and CaDPA release at single spore level. However, the biochemical detail for the trigger signal or germination substance P remains unclear.

In the current work, we have attempted to develop a mathematical model to identify a minimal reconstructed molecular network that is crucial for germination kinetics and memory. We consider a minimal set of GRs in two states – inactive and active, and a SpoVA channel in three states - closed inactive, closed active and open. When the number of SpoVA proteins in the open active state reaches a threshold level (e.g. 50 copies/spore used in this paper), the channel is largely opened for rapid CaDPA release. Numerical simulation reveals that memory can reside with GRs and/or be stored in the CaDPA channel complex, which is consistent with the previous proposal that spore memory is stored in metastable states of SpoVA proteins (32). Obviously, this model must be able to predict both the heterogeneity of *T*_lag_ and the percentages of spore germination in 1^st^ and 2^nd^ germinant pulses. In addition, with computer simulations we can identify network topologies that exhibit minimal deviation between predicted and experimental data, and reveal reaction dynamics of molecular processes between GR activation and rapid CaDPA release. Finally, analysis of the optimized model could shed light on the question of how the network’s molecules and their interactions determine its biological function.

## Results

### Spore germination model

In our modelling strategy, we follow a top-down approach. First we develop an overall mathematical model and then we dissect the mathematics into a set of mass action kinetics based differential equations based on the components of the germinosome and the SpoVA channel which must somehow interact. As a boundary condition we opted for the most parsimonious number of building blocks, and that the model must be able to recreate published experimental spore memory data (32). One common premise for commitment in molecular networks is a bistable switch (37). In such bistable systems two stable states coexist, and thus spores are dormant or are committed to germinate (38). Upon activation, the system is irreversibly committed to one state. A condition for bistability is non-linear dynamics involving positive feedback in which a molecular species upregulates its production or autoactivates its function, *e.g.* channel opening, a mechanism to filter out noise and a mechanism to prevent numeric explosion of the upregulated or activated species (38,39). A simple deterministic molecular kinetic model for the autoactivation and deactivation rates of the state variable *X* commonly used to model bistability is given in Eq (1) (40,41).

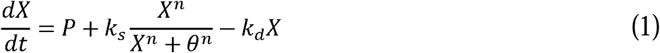

In this equation, *X* denotes the concentration [molecules/spore] of a crucial state of a germination protein, *k*_s_ is its maximum production rate, *n* represents cooperativity of variable *X*, *θ* ^n^ sets the value at which the rate of production of *X* is half its maximum value, *k*_d_ is the decay rate of protein *X* and *P* is a basal production rate (40–42). Positive feedback is introduced by cooperative action of the system variable *X* for *n* >1. The multistabilty of Eq (1) depends on the settings of the various parameters. There are some settings in which *X* gradually increases upon an inductive stimulus and there is a parameter subspace where *X* exhibits a bistable response. In the latter case, there are two stable states, one at low *X* and one at high *X*, and an unstable state in between. Fluctuations of *X* mediated by *P* around the lower stable state (state 1) but not progressing to the metastable state fall back to the lower stable state. Fluctuations of *X* through *P* surpassing the unstable state are auto-activated and increase to the stable state 2 at high *X*. Fluctuations in *X* above the high stable state fall back to stable state 2, thereby preventing a runaway of variable *X* to infinity. Eq (1) is thus an autocatalytic off-on switch with parameter subspace for bistability and is the core of our model.

Starting from this equation we designed an input section, signal transduction and finally we describe SpoVA channel opening kinetics to estimate *T*_lag_ times in germination. The molecular wiring of the reconstructed molecular network of the model is given in Fig 1. In our model, we do not differentiate between GerA, GerB and GerK GRs, and thus do not differentiate between germinants. GRs can be in an inactive *R*_i_ state or an active *R*_a_ state, and germinant *S* binding converts an *R_i_* GR to *R_a_* with a rate constant *k_1_*. Low copy protein numbers in individual cells in bacterial populations usually vary according to a gamma distribution (43). Measurements of levels of GFP-labelled GRs in spores have shown that GR copy numbers also vary according to a gamma distribution function (43,44). Thus, the distribution of total levels of *R*_i_ in spores in the absence of germinant is given by the probability density (gamma) function

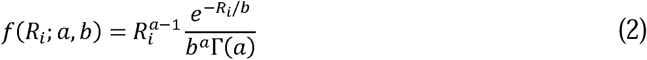

in which Γ(a) = (a-1)!, *a* is the shape parameter and *b* is an inverse scale parameter. The distribution mean *E*(*R*_i_) = *ab* and the variance *Var*(*R*_i_) of the distributions is *ab*^2^. The skewness of the distribution is equal to 2/*a*^½^. The skewness depends only on the shape parameter *a* and approaches a normal distribution when approximately *a* > 10.

**Fig 1.**
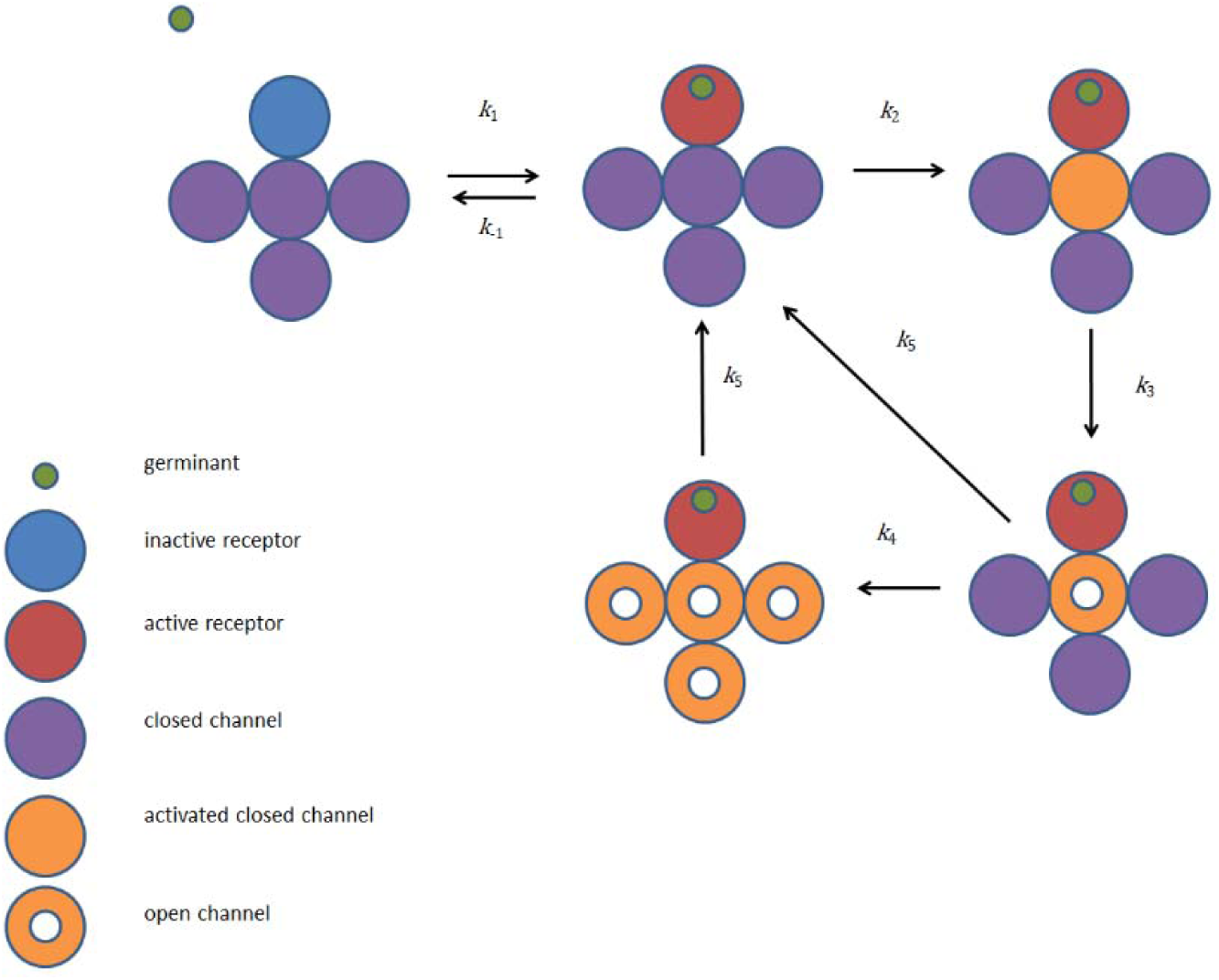
Architecture of a *Bacillus subtilis* germinosome that exhibits spore germination memory. The molecular network is composed of GR proteins that are activated by a germinant, and channel proteins are in a closed, activated closed and an open state and undergo mutual interactions.

Germinant binding in our model is reversible and dissociation takes place with rate constant *k*_-1_. Eq (3) and Eq (4) describe rates of formation of active GR molecules *R_a_*and rates of depletion of the inactive GR population *R_i_*. In contrast to GRs which we propose exist in only two states, *R*_i_ and *R*_a_, we need a SpoVA channel that can adopt three states. While there is no experimental proof for the latter assumption, a model with two SpoVA states does not reproduce experimental germination curves (data not shown). The SpoVA channel *C*_c_ is closed and inactive, the channel *C_ca_*is closed and in an activated state, or the channel *C_o_* is open and in an activated state. Interaction of activated GRs *R_a_* with closed SpoVA channels *C_c_* activates the latter (Eq (5)). We assume a direct interaction, although there is no conclusive experimental data about signaling from activated GRs to the SpoVA channel. However, yeast two hybrid experiments suggest that there is interaction between *B. subtilis* GerA proteins and several SpoVA proteins (45). The function of GerD is ignored in the model. A premise for commitment in molecular networks is a bistable switch with delay. Bistability is introduced here through intersection of a production function comprising a Hill function with *n* > 1 and a linear decay function in Eq (7). The production function describes the rate of opening of SpoVA channels. Cooperativity of open SpoVA channels also contributes to the opening rate. This means that open SpoVA channels (C*_o_*) have multiple mutual interactions which enhance the opening rate of closed channels. Cooperativity of SpoVA channel gating is introduced through a Hill function with apparent dissociation constant *θ* and coefficient *n*. The decay function describes closure of channels with reaction rate –*k*_5_*C_o_*. Channel closure is first order in the concentration of open channels. Upon closure the closed channel falls back to the inactive ground state *C*_c_.

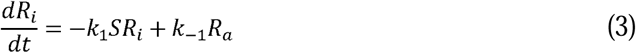

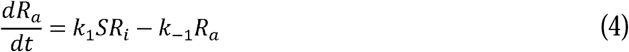

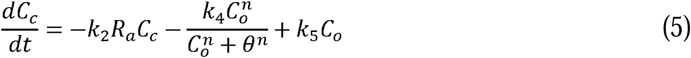

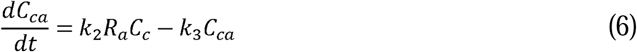

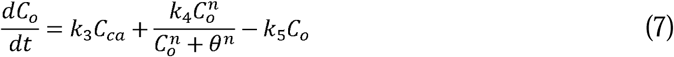

In these equations, *S* is germinant concentration, *R_i_*inactive GR, *R_a_* active GR, *C_c_* closed SpoVA channel, *C_ca_* activated closed SpoVA channel, and *C_o_* open SpoVA channel. The units of the GR and SpoVA variables are numbers of the various complexes/spore. Constants *k*_1_, *k*_-1_, *k*_2_, *k*_3_, *k*_4_ and *k*_5_ are rate constants, *n* is a Hill coefficient, and *θ* ^n^ sets the value at which the production rate in the second term is half its maximum value. The final stage of the model is SpoVA channel opening. Whether core CaDPA is exported by concentration differences and/or an electrochemical gradient of ions inside and outside the inner membrane, CaDPA efflux will be proportional to the number of open SpoVA channels, and we assume that *T*_lag_ is determined by a minimal number of open SpoVA channels needed for rapid CaDPA release. It is known that CaDPA activates the CLE CwlJ and this enzyme’s action stimulates CaDPA efflux. However, CwlJ activation and its effect on the SpoVA channel opening rate is not yet a part of the model, and we expect that this effect will become significant only after *T*_lag_. If there is a contribution of CwlJ activation to SpoVA opening than this will lead to a minor shift of *T*_lag_. In our computer model, we use a cycle of 500 numeric simulations to solve the set of coupled ordinary differential equations. In each simulation, we predict the number of active open SpoVA channels as a function of time and we determine for each simulation the time of opening, *T*_open_, of SpoVA channels. This time point corresponds with the time point at the beginning of fast CaDPA release (28), and is related to *T*_lag_ through *T*_lag_ = *T_open_* – *T*_block_, where *T*_block_ is the time of the end of a germinant pulse. We define *T*_open_ as the time point in a simulation where *C*_o_ exceeds 50 copies/spore. This number is arbitrarily selected as experimental data are lacking.

### Modeling of spore memory of germinant stimuli

First, we reproduce with our mathematical model an earlier published percentage germination curve showing spore memory for stimulation of wild-type *B. subtilis* spores with L-valine (32). Next, we study the relationship between the mean *E*(*R*_i_) of the GR distribution and the percentage germination upon the 1^st^ and the 2^nd^ germinant pulses. We then compare the predictions with experimental results of the percentage germination of wild-type spores and mutant spores with elevated GR levels. Wild-type spores can acquire memory upon short pulses of L-valine to induce germination as follows. After the 1^st^ pulse a fraction of the spores is committed to germinate and does so. From the subpopulation that does not germinate after the 1^st^ pulse, a fraction germinates upon a 2^nd^ pulse and finally a fraction does not germinate at all. Interestingly, spore populations can exhibit 2- to 3-fold more germination in the 2^nd^ germinant pulse following the 1^st^ one. Apparently, there is memory in the molecular system for external perturbations such as a germinant pulse that occurred before the 2^nd^ pulse (32). Consequently, we optimize model parameters to reproduce the spore germination curve for an experiment with two short germinant pulses (32). In our simulation, we select two pulses of germinant exposure from 0 < *t* < 5 min and 30 < *t* < 35 min. We set the germinant concentration to 3.5 mM. A percentage germination curve in our definition is a cumulative distribution function of *T*_open_ to give rapid CaDPA release. Spores that do not germinate have an infinite *T*_open_ and do not contribute to the distribution in the time frame of the simulations. Heterogeneity of *T*_open_ and as result *T*_lag_ determines the shape of the curve. Here, we introduce heterogeneity through a gamma distribution of GR copy numbers (Eq (2)). For each simulation of a cycle of 500 we randomly select a GR copy number from a gamma distribution, but keep SpoVA copy numbers constant. This means that *C*_c_ + *C*_ca_ + *C*_o_ is constant in each cycle of a set of simulations. This approach is justified by the experimental observation that overexpression or deletion of GR’s has no effect on levels of SpoVAD and by inference levels of all proteins encoded by the heptacistronic *spoVA* operon (15). From Western blotting, it is known that on average ∼700 to ∼1,100 of each GR are present in a spores’ germinosome. The starting value of at least SpoVAD is 6,500 as determined earlier by Western Blotting (20). By tuning of *k*_4_, the SpoVA copy number of activated and open channels in our simulations asymptotically approaches 6,500 copies/spore (*vide infra*). This copy number is also an arbitrary selection as we do not know by experimentation the number of open SpoVA channels at the end of rapid CaDPA release. At the end of the simulations we construct a predicted percentage germination curve. The experimental germination curve was determined by monitoring germinating spores with DIC microscopy. However, most of the intensity change seen in DIC microscopy is caused by rapid CaDPA release, and the kinetic parameters for which averages were determined are *T*_lag_, *T*_release_ and Δ*T*_release_ = *T*_release_-*T*_lag_ (28,32,46).

We start with varying the mean *E*(*R*_i_) of the GR distribution and keep the standard deviation *Var*^½^(R) fixed. The parameters of the randomly accessed gamma distribution are estimated at the end of a set of 500 cycles from the fitted gamma distribution. A typical GR protein distribution is depicted in Fig 2A. The average of this GR copy number distribution is set to 1100 copies/spore and we selected combinations of distribution parameters *a* and *b* with the condition that *Var*^½^(x) = *a^½^b* = 220. With this setting the GR distribution tails to higher copy numbers, and a fraction of the spores germinate after the 1^st^ pulse and a fraction germinates after the 2^nd^ germinant pulse (Fig 2A, B). Notably, with the 1^st^ pulse 15% of the spore population germinates, while after the 2^nd^ pulse 68% of the remaining spores germinate. Although the germinant concentration and pulse lengths are equal for both pulses, germination after the 2^nd^ pulse is approximately four-fold higher than after the 1^st^ pulse. Thus, the model reproduces published experimental data (32). Simulations for other *E*(*R*_i_) are compiled in S1 Figures.

**Fig 2.**
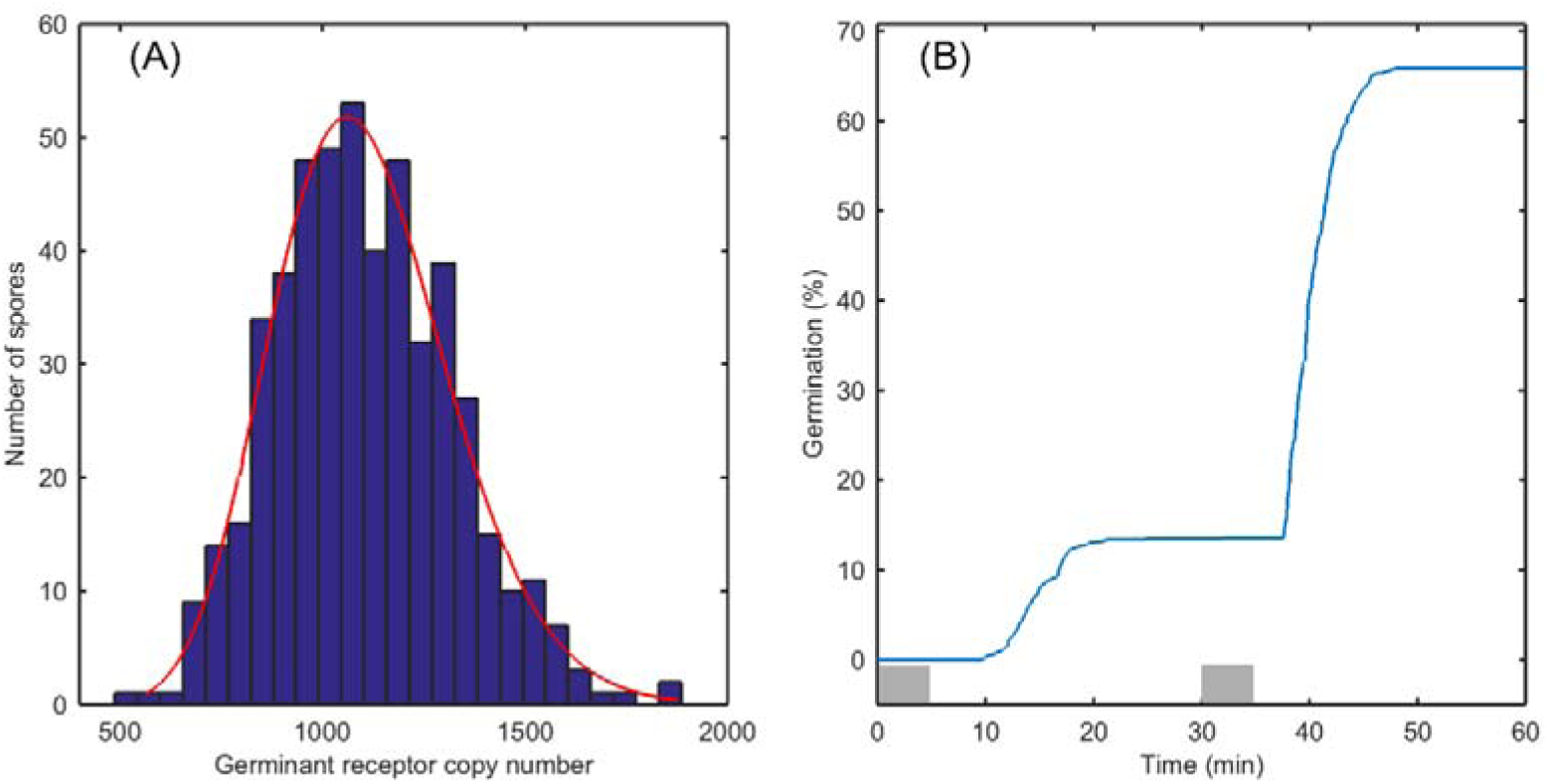
Simulation of spore germination memory (A) Gamma distribution of GR copy numbers used for simulation of the percentage germination efficiency curve (Fig. 2b) with *E*(*R*_i_) = 1100 at *t* is 0 min. The red line is the fitted distribution. The GR estimated mean is 1105 molecules/spore, and the estimated standard deviation is 219 molecules/spore; the parameters of the fitted gamma distribution are: *a* is 25.4491 [22.5002, 28.7844] and *b* is 43.4004 [38.3247, 49.1484]; Right panel: Percentage germination of spores in a two pulse germination experiment. (B) Percentage germination in a two pulse germination experiment. After the 1^st^ pulse the curve levels off to an asymptote at 15%. After the 2^nd^ pulse the curve approaches 68%. Two pulses of 3.5 mM germinant for receptor *R*_i_ activation are administered from 0 < t < 5 and 30 < t < 35 min. At zero time *R*_a_ = 0 copies/spore; *C*_c_ = 6500 copies/spore. *C*_a_ = *C*_o_ = 0 copies/spore; *k*_1_ = 14e-3; *k*_-1_ = 100e-3; *k_2_* = 25e-7; *k*_3_ = 60e-3; *k*_4_ = 195e2; *k*_5_ = 305e-2; *n* = 3; *θ* = 20.

We repeated a double pulse germination experiment with *B. subtilis* wild-type spores and spores of an isogenic strain that overexpresses the GerA GR ∼8-fold as determined by Western Blotting, with an experimental error of 25% (7). The experimental percentage germination curves are depicted in Fig 3A, B. We simulated the curves with our model (Fig 3C, D). Compared to germination of wild-type spores we had to adjust reaction rate constants for a 2 min pulse for wild-type spores that overexpress GerA with 10 mM L-valine as germinant. A broad GR copy number distribution, with *E*(*R*_i_) at 1100, *a* is 1 and *b* is 1100, reproduces approximately 4% germination upon the first pulse and approximately 6% germination at the second pulse (Fig 3C). Next, we overexpress the GerA GR by increasing *E*(*R*_i_) and keep the reaction rate constants similar. In this approach, we put forward the constraint that overexpression of GR GerA from a more active promotor does not affect the rate constants for binding and dissociation of the germinant to the receptor. A second assumption is that expression of GerA from another promotor affects the parameters of the gamma distribution. It is known from modelling of burst like stochastic gene expression that the rate at which the gene transitions from the inactive state to the active state and vice versa determines the momenta of the mRNA copy number distribution and consequently the protein copy number distribution (47). Our modelling results are in accord with an earlier prediction using a stochastic gene expression model that a high burst frequency produces a narrow mRNA copy number distribution and a low burst frequency a broad mRNA distribution (47). However, experimental data for the distribution of copy numbers of an overexpressed GFP-labelled GerA in individual spores are lacking. With *E*(*R*_i_) at 3350 *a* as 67 and *b* as 50 we reproduce 3% germination at the first pulse and 64% germination at the second pulse (Fig 3D). In this case, we had to adjust the variance of the GR distribution to reproduce the percentage germination curve. For *E*(*R*_i_) at 3350 the overexpression is approximately 3.3-fold. For further increase of *E*(*R*_i_) we have to diminish *k*_1_ to keep the second pulse at 63% without surpassing the 3% germination at the first pulse. These results validate simulations of the relation between GR protein distributions and percentage germination curves in a double pulsed memory experiment. Interestingly, the model predicts constant Δ*T*_release_ times and that Δ*T*_release_ is independent of *T*_lag_, corroborating observations from earlier experimental work (28). Next, we probed for the effect of varying the concentration of germinants used during the pulses, from 2 mM to 5 mM (S2 Figures). At 2 mM germinant no germination is predicted with the 1^st^ pulse and approximately 1% after the 2^nd^ pulse. This latter percentage increases to 12% at 2.5 mM germinant, but still with no germination after the 1^st^ pulse. This implies that the 1^st^ pulse primes 12% of the spore population for germination after the 2^nd^ pulse. For higher germinant concentrations, significant germination with the 1^st^ pulse as well as substantial memory effects are predicted, *e.g.* S2 Fig 2C. The influence of the standard deviation of the distribution is further highlighted in S3 Figures. There is an obvious relationship between the variance of the distribution and the fraction of spores germinating after the 1^st^ and 2^nd^ pulses (S3 Figures). As the variance of the distribution increases and tails to higher GR copy numbers, more spores germinate with the 1^st^ germinant pulse.

**Fig 3.**
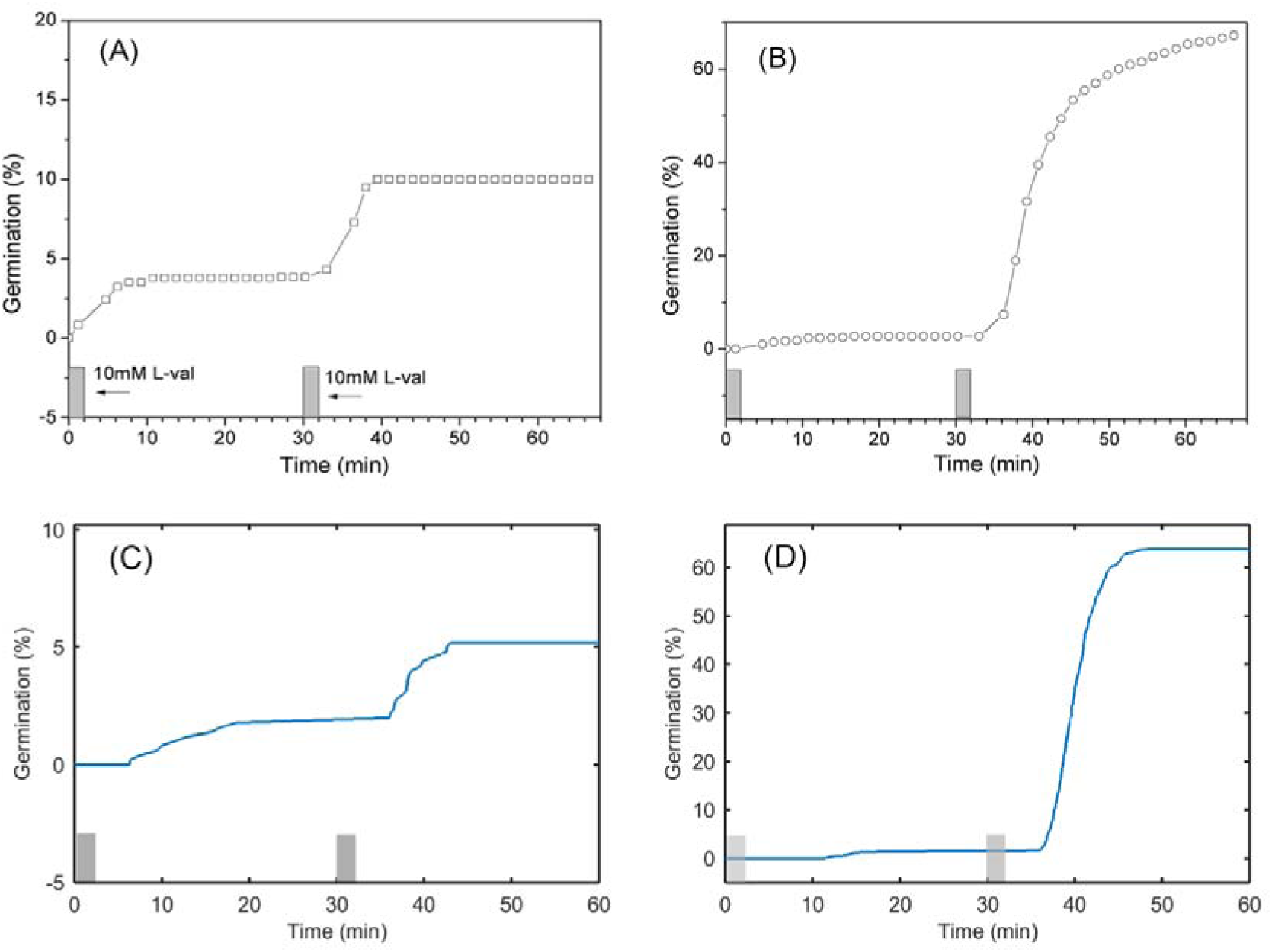
A double pulse germination experiment with *B. subtilis* wild-type spores and spores of an isogenic strain that overexpresses the GerA GR. (A) Wild-type *B. subtilis* spores (PS533) were given two 2-min 10 mM L-valine (L-val) germinant pulses separated by 30 min. (B) Spores of *B. subtilis* PS3476 (GerA GR over-expressed ∼ 8-fold) were given two 2-min 10 mM L-valine germinant pulses separated by 30 min. Gray bars above the horizontal axis indicate pulse durations. (C) Simulated percentage germination of WT spores in a two pulse germination experiment. The estimated distribution mean is 1118 and the standard deviation is 1064 with *a* is 1.10476 [0.989585, 1.23334] and *b* is 1012.19 [881.615, 1162.1]. At zero time *R*_a_ = 0 copies/spore; *C*_c_ = 6500 copies/spore. *C*_a_ = *C*_o_ = 0 copies/spore; *k*_1_ = 350e-5; *k*_-1_ = 100e-3; *k_2_* = 25e-7; *k*_3_ = 60e-3; *k*_4_ = 195e2; *k*_5_ = 305e-2; *n* = 3; *θ* = 20. (D) Simulated percentage germination of overexpressed GR spores in a two pulse germination experiment. The estimated distribution mean is 3367 and the standard deviation is 399 with *a* is 71.2149 [62.9306, 80.5898] and *b* is 47.2855 [41.7667, 53.5336]. At zero time *R*_a_ = 0 copies/spore; *C*_c_ = 6500 copies/spore. *C*_a_ = *C*_o_ = 0 copies/spore; *k*_1_ = 350e-5; *k*_-1_ = 100e-3; *k_2_* = 25e-7; *k*_3_ = 60e-3; *k*_4_ = 195e2; *k*_5_ = 305e-2; *n* = 3; *θ* = 20.

### Dynamics of SpoVA channel opening

The kinetics of SpoVA channel opening are studied in more detail for GR copy numbers ranging from 600 to 1,800 molecules/spore. The increment of GRs is 50 copies per cycle of numeric evaluation of the set of ordinary differential equations (ODE’s) in a simulation. The number of GRs is defined by *R*_i_ and at time point *t* is 0 seconds. The results *C*_o_ and the copy number time profiles of the state variables *R*_i_, *R*_a_, *C*_c_, and *C*_ca_ are given in Fig 4. There are three subpopulations in the uniform GR distribution. With 600 to 950 GRs/spore, there is no germination, with 1000 to 1350 GRs/spore there is germination after the 2^nd^ pulse and with 1400 to 1800 GRs/spore there is germination after the 1^st^ and 2^nd^ pulses. Fig 4 shows that only in a fraction of the spore population are GRs sufficiently activated by germinant binding to lead to commitment to germinate. We can tune this fraction by variation of the rate constants *k*_1_ and *k_-_*_1_ for binding of *S* to GR and dissociation thereof. This tuning also affects the number of spores opening SpoVA channels during the 1^st^ and 2^nd^ pulses. With the parameter settings of our simulation only a few SpoVA channels are activated by interaction with GRs. Because of different copy numbers between GRs and channels there cannot be a one to one stoichiometry between GR activation and channel opening. Approximately 1,100 GerA GRs are in the germinosome, while ∼6,500 SpoVA channels are dispersed over the IM. The surface area of the spore IM can be approximated by an ellipsoid. The surface area of an ellipsoid (*A*_im_) is calculated as *A*_im_ = 2π(*s*^2^+*ls*), where *l* is the length of the long axis and *s* of the short axis (48). The long axis *l* and short axis *s* of the IM are estimated by electron microscopy and *l* is 837 nm and *s* is 554 nm for a typical spore (49). These are obtained by subtraction of coat and cortex layers’ thickness from the spore long axis and short axis (49). The surface area of the IM is *A*_im_ = ∼ 4,835,800 nm^2^, and on average there will be one SpoVA molecule per 744 nm^2^ of IM. Next, we calculate the area of the germinosome. As input we use the normalized average distribution of the fluorescence signal along the long axis of a spores. The full width at half max intensity for GerKB-GFP signal is 300 nm (9). Assuming a planar circle on the surface of the IM for the contour of the germinosome its surface *A*_ger_ = π*r*^2^ = 502,655 nm^2^. Assuming a homogeneous distribution of 744 nm^2^ per SpoVAE then the germinosome contains at most 676 SpoVA molecules. In a stoichiometry where one GR activates one channel maximally, 1,100 channels can be activated. Thus, there are ∼ 5,800 SpoVA channels that cannot directly interact with a GR. Thus, SpoVA channel cooperativity is introduced in the model to relay the signal from GRs over the entire IM surface to open more than approximately 676 channels. In our model, closed SpoVA channels are directly opened by activated and open SpoVA channels. Immediately after the 1^st^ germinant pulse the number of open SpoVA channels gradually increases. This slow increase is the gross difference between opening by activated SpoVA channels through *k*_4_*C*_a_ and closing of SpoVA channels through decay –*k*_5_*C*_o_ (Eq (7)). Beyond 10 min sufficient channel opening takes place to force cooperative opening of the dark blue subpopulation through the Hill function term in Eq (5). Channel opening is autocatalytically driven, all SpoVA channels open, and hence the *C_o_*curve approaches 6,500 molecules. The SpoVA channels from spores that germinate after the 2^nd^ pulse are partially activated by the 1^st^ pulse but remain below a critical copy number needed to force an autocatalytic opening mediated by the Hill function in Equation 6. However, the extent of channel opening does not fully return to the baseline. A 2^nd^ pulse at 30 < *t* < 35 min contributes additionally to channel opening and drives through cooperative action channel opening to completion at 6,500 copies/spore. As a result, channels open that are not opened by the 1^st^ germinant pulse. Thus, the system exhibits memory for an earlier germination stimulus. Finally, the curve levels off to a plateau at 6500 open SpoVA channels. The level of the plateau of open SpoVA channels is mainly determined by *k*_4_ and *k*_5_. A third fraction does not open at all, as the 2^nd^ germinant pulse is insufficient to trigger autocatalytic SpoVA opening and the number of open channels decays to the baseline.

**Fig 4.**
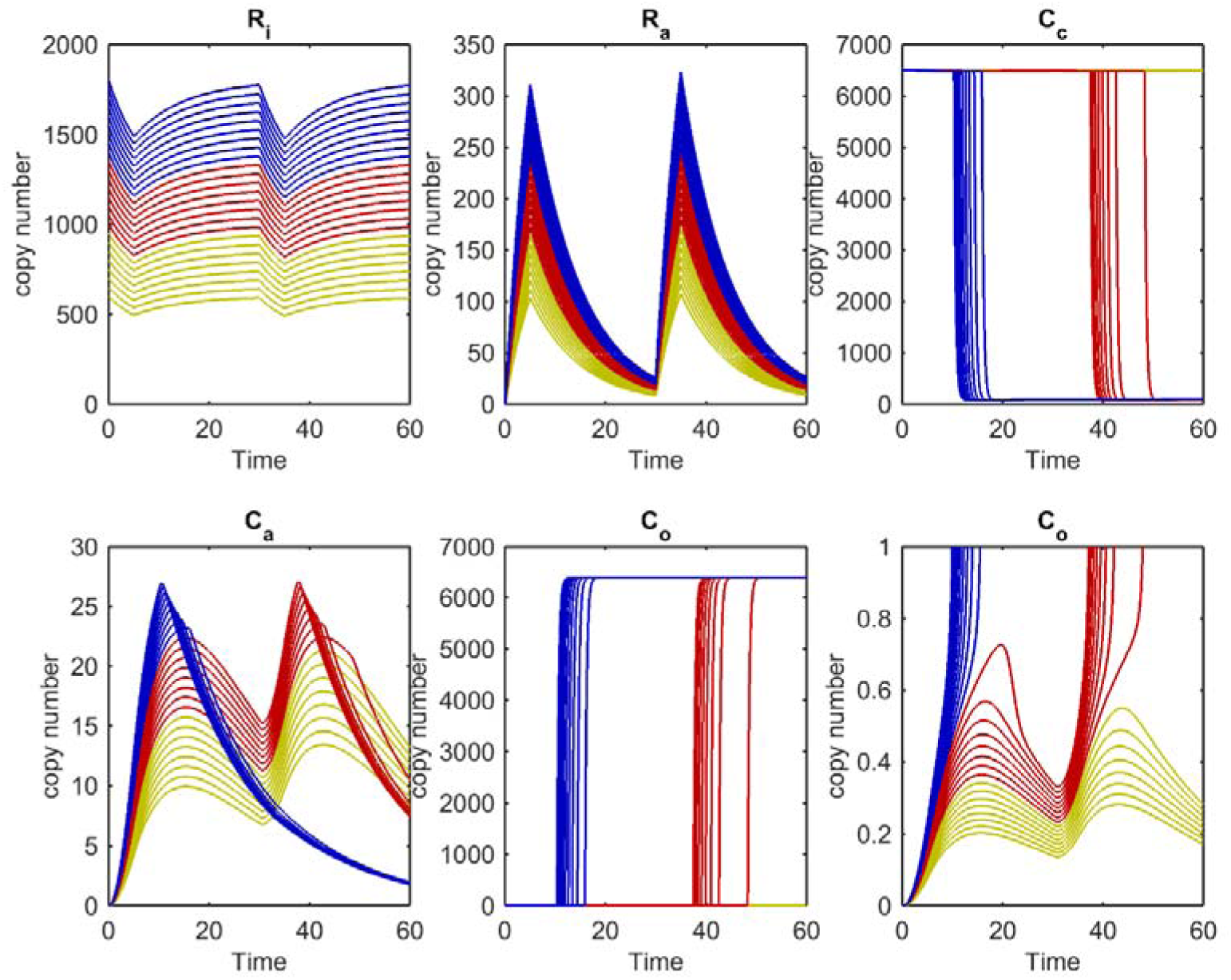
The copy numbers of *R*_i_, *R*_a_, *C*_c_, *C*_a_, *C*_o_ and an extension of *C*_o_ as a function of time. Each trace represents a state variable profile for a variable GR *R*_i_ copy number. The starting GR values are 600 ≤ *R*_i_ ≤ 1800 copies/spore and the increment is Δ*R*_i_ = 50 copies/spore. The yellow spore population (600 ≤ *R*_i_ ≤ 950) does not germinate, the red one (1000 ≤ *R*_i_ ≤ 1350) germinates at the 2^nd^ pulse and the blue ones (1400 ≤ *R*_i_ ≤ 1600) germinate at the first pulse. Two pulses of 3.5 mM germinant for *R*_i_ activation are administered from 0 < t < 5 and 30 < t < 35 min. At time zero, *R*_a_ = 0 copies/spore; *C*_c_ = 6500 copies/spore. *C*_a_ = *C*_o_ = 0 copies/spore; *k*_1_ = 14e-3; *k*_-1_ = 100e-3; *k_2_* = 25e-7; *k*_3_ = 60e-3; *k*_4_ = 195e2; *k*_5_ = 305e-2; *n* = 3; *θ* = 20.

## Discussion

Spore germination experiments with sequential germinant stimuli separated in time are very suitable to construct and optimize a kinetic germination model. The shape of the percentage germination curves with multiple inflection points and asymptotes enables a more robust fitting of model parameters than a single stimulus sigma-shaped germination response experiment. The germination model based on the minimal molecular network as presented in Fig 1 and described by differential Eq (3–7) reproduces percentage germination curves in double pulse memory experiments (32) and newly acquired data (Fig 3A, B). Upon germinant exposure in two 5 min pulses separated by thirty min, two subpopulations of the whole population germinate. The second one exhibits a memory effect, in that the amounts of spores that germinate after the 2^nd^ pulse are larger than those germinating after the first pulse. Memory has been attributed to a conformational change in some germination protein, that activates this protein for germination, although the activated protein can also decay to the inactivate ground state. When the activated protein is not fully decayed, then a 2^nd^ germinant pulse leads to a larger population of activated proteins and a higher percentage of germination after the 2^nd^ pulse (32). The reconstructed network shows that a minimal set of an inactive and active GR and a transport channel in three states *-* closed inactive, closed active and open is needed to reproduce memory. Thus, memory can reside with GRs and/or be stored in the CaDPA channel complex. This is consistent with generation of memory in spores that lack all GRs when germination is triggered by direct or indirect stimulation of CaDPA release by exogenous CaDPA itself that activates CwlJ, or by dodecylamine that may cause the SpoVA channels to open (32).

Memory is stored in an activated GR *R*_a_, although this can decay to the inactive state *R*_i_. As is seen in Fig 4, GR inactivation does not fully return to the baseline before the 2^nd^ germinant pulse and thus GR inactivation characteristics contribute to memory. The fraction of activated GRs and its relaxation to an inactive state is determined in our model by the ratio *k*_1_/*k*_-1_, although we do not know this ratio from experimental data. However, memory in our model is not uniquely confined to GR activation and inactivation kinetics, as there is a model parameter setting with *k*_-1_ = 225.10^-3^ and *k*_2_ = 45.10^-7^ where *R*_a_ returns to baseline before the 2^nd^ germinant pulse and the network still shows memory for germination without adjusting the other rate constants. In this case memory is introduced in the network by the interplay between activation of closed channels and their rate of opening to *C*_o_. The spore channel activation rate is proportional to *k*_2_*R*_a_ and the number of closed inactive SpoVA channels *C*_c_ (Equation 6), and the rate of opening is proportional to *k*_3_*C*_a_. We assume that the number of activated GRs at 3.5 mM is approximately 17% and we have optimized *k*_2_ accordingly. This needs further experimentation to dissect *k*_2_*R*_a_ into *k*_2_ and *R*_a_. Rate constant *k*_3_ has an effect on *T*_lag_ because it feeds the rate of open channel formation (Eq (7)). The Hill coefficient *n*, *θ* together with *k*_4_ and *k_5_* determine the bistability of the model needed to irreversibly open CaDPA channels. In this simulation *n* is 3 pointing to cooperativity effects of open activated SpoVA channels (Eq (7)). Furthermore, rate constants *k*_4_ and *k*_5_ in combination with *θ* determine the number of open SpoVA channels at the end- point of simulations. This number is not known by experimentation and is arbitrarily set.

The time to the onset of rapid CaDPA release *T*_lag_ is heterogeneous and shows a probability distribution. We introduce germination time heterogeneity through the GR distribution in spore population. In a cycle of computer simulations, we keep the rate constants fixed. Rate constants *k*_1_ and *k*_-1_ are on-rate and off-rate constants and determined by the 3-D structure of the GR proteins. As the primary sequence of the GR proteins is not variable within an isogenic strain, we expect that a *k*_1_ and *k*_-1_ probability distribution does not contribute to heterogeneity of *T*_lag_ at the population level. At present, our model is a minimal one, and we do not differentiate between actions of GerA, GerB and GerK GRs. Consequently, we do not integrate the signal from different GRs and germinants. We also do not know the fraction of SpoVA channels in an inactive state that is activated by a population of activated GRs. The network in our model is capable of signal amplification, as with current parameter settings approximately 1-27 closed SpoVA channels are activated (Fig 4). This is sufficient to irreversibly open the entire channel population through cooperative opening which is needed to open channels outside the perimeter of the germinosome. Another benefit from the network wherein only fractions of molecular populations are active is that it dampens fluctuations in germinant exposure. This ensures that there is a threshold for GR activation that will prevent unwanted early germination.

The model generated in this work has been optimized to reproduce memory of germinant exposures. Other combinations of reaction rate constants will likely also give kinetics that show memory, but the molecular wiring and the kinetic differential equations will likely remain identical. However, further, experimentation is needed to narrow down the parameter space. In particular, the contributions of GR activation and deactivation, and SpoVA channel activation, deactivation and opening should be studied. The endpoint of germination in our model is the rapid opening of SpoVA channels. In a future extension of the model, the effects of the CLEs SleB and CwlJ should also be included. In addition, as noted above GR-independent germination by CaDPA or dodecylamine also show memory, presumably through effects on the SpoVA channel alone, and this can also be introduced into the model. The model in its current state will be used in future research to reduce the number of experimental variables in an iterative cycle of prediction of the dynamics of a hypothetical molecular network, the design of an experiment, comparison with the outcome, and further tuning of the model until agreement between theoretical predictions and experimental data is reached.

## Materials and methods

Simulations were carried out using the MATLAB package of software (the Mathworks, Natick, USA). M-files with source code for simulation of percentage germination curves and copy number time profile plots are given in S Matlab m-file one and S Matlab m-file two.

### Phase contrast microscopy

Spores (1 μl; ∼10^8^ spores/ml in water) were spread on the surface of a microscope coverslip that was then dried in a vacuum desiccator for 5 min. Coverslips were suspended in water and then mounted on and sealed to a microscope sample holder kept at a constant temperature of 37°C. A phase contrast microscope was used to record the images of ∼500 individual spores at a rate of 15 s per frame. Spores that contain DPA appear as bright and spores that do not contain DPA appear as dark. Percentage of spores containing DPA was defined as the number of bright spores divided by the total number of spores.

### Spore germination with germinant pulses

*B. subtilis* spores of strains PS533 (wild-type) and PS3476 (overexpresses the GerA GR ∼ 8-fold) were prepared, purified and stored as described previously (14). The purified spores were free (>98%) of sporulating or growing cells or germinated spores as observed by phase contrast microscopy. *B. subtilis* spores were heat activated prior to nutrient germination by incubation in water at 70°C for 30 min and then by cooling on ice for at least 15 min. *B. subtilis* spores attached on the surface of a glass coverslip were germinated in 25 mM K-HEPES buffer (pH 7.4) with a pulse of a germinant (10 mM L-valine) for 2 or 4 min before the germinant was removed and spores were rinsed 5 times with 25 mM HEPES buffer (pH 7.4) using vacuum pump suction (32). The rinsed spores were then incubated at 37°C in 25 mM K-HEPES buffer with a pH of 7.4 unless noted otherwise. After various incubation times, the spores were given a 2^nd^ exposure to germinants for various times at 37°C in 25 mM K-HEPES buffer (pH 7.4) followed by germinant removal, rinsing by vacuum pump suction, and further incubation at 37°C in 25 mM K-HEPES buffer (pH 7.4).

## Acknowledgements

CdK, LdK and SB acknowledge the University of Amsterdam for funding of their research.

## Author contributions

CdK developed the model and performed computational analyses with input from LdK and SB. XL and YQL conducted experiments. PS provided spores of *Bacillus subtilis* strains. CdK wrote the manuscript with input from LdK, SB, YQL and PS. YQL, JL and PS designed the germination experiments.

## Conflict of interest

The authors declare that they have no conflict of interest.

## Notes

### Competing Interest Statement

The authors have declared no competing interest.

### Summary of Updates

New version of the main text and supplementary files

