## Supplementay Figure S3 for "A mathematical kinetic model of memory in *Bacillus subtilis* spore germination"

Supplementary Material 3

The influence of GR gamma distribution standard deviation on percent germination in the 1^st^ and the 2^nd^ germinant pulses. The scale and rate parameters *a* and *b* are varied where at *t* is 0 min the mean GR copy number is *E*(*R*_i_) = 1100 copies/spore and the standard deviation *Var*^½^(*R*_i_) is 47, 105, 348, 778 and 1100 copies/spore. The parameters and initial values of remaining variables at time zero are *R*_a_ = 0 copies/spore; *C*_c_ = 6500 copies/spore. *C*_a_ = *C*_o_ = 0 copy/spore; *k*_1_ = 14e-3; *k*_-1_ = 100e-3; *k_2_* = 25e-7; *k*_3_ = 60e-3; *k*_4_ = 195e2; *k*_5_ = 305e-2; *n* = 3; *θ* = 20. The germinant receptor protein copy numbers are randomly drawn from a Gamma distribution.

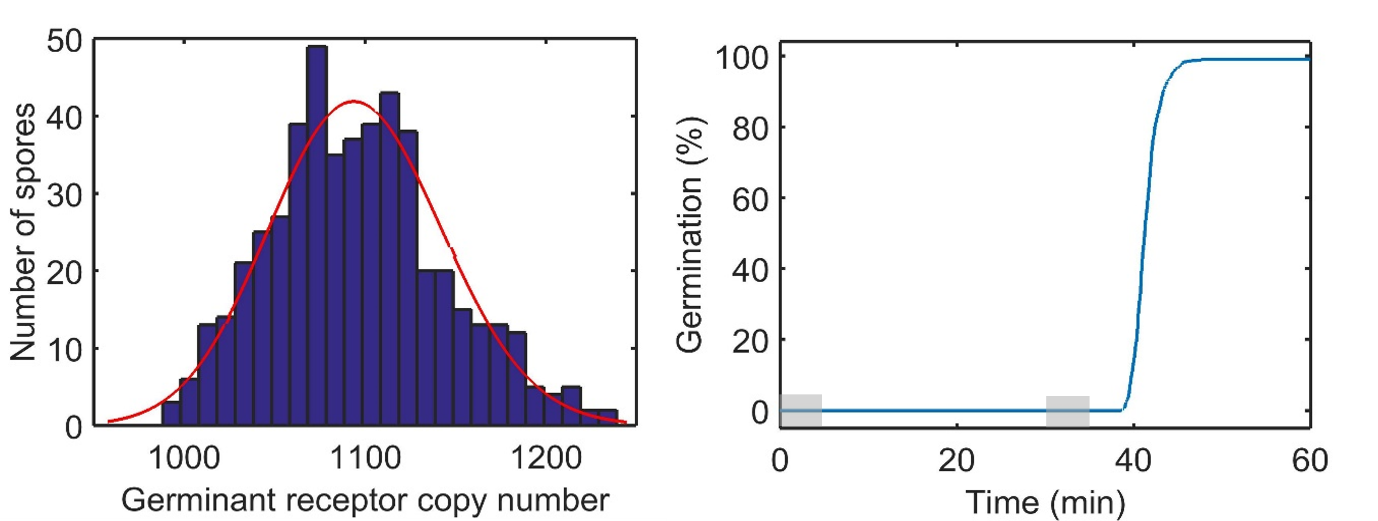

Fig S3a.	Left Panel: Gamma distribution of GR copy numbers used for simulation of the percentage germination efficiency curve (Right panel). The red line is the fitted distribution. *Var*^½^(*R*_i_) is 47 at *t* is 0 min. The GR estimated mean is 1096 copies/spore, and the GR estimated standard deviation is 48 copies/spore; the parameters of the fitted gamma distribution are: *a* is 524.484 [463.356, 593.676] and *b* is 2.08941 [1.84578, 2.36519]. Right panel: Percentage germination of spores in a two pulse germination experiment.

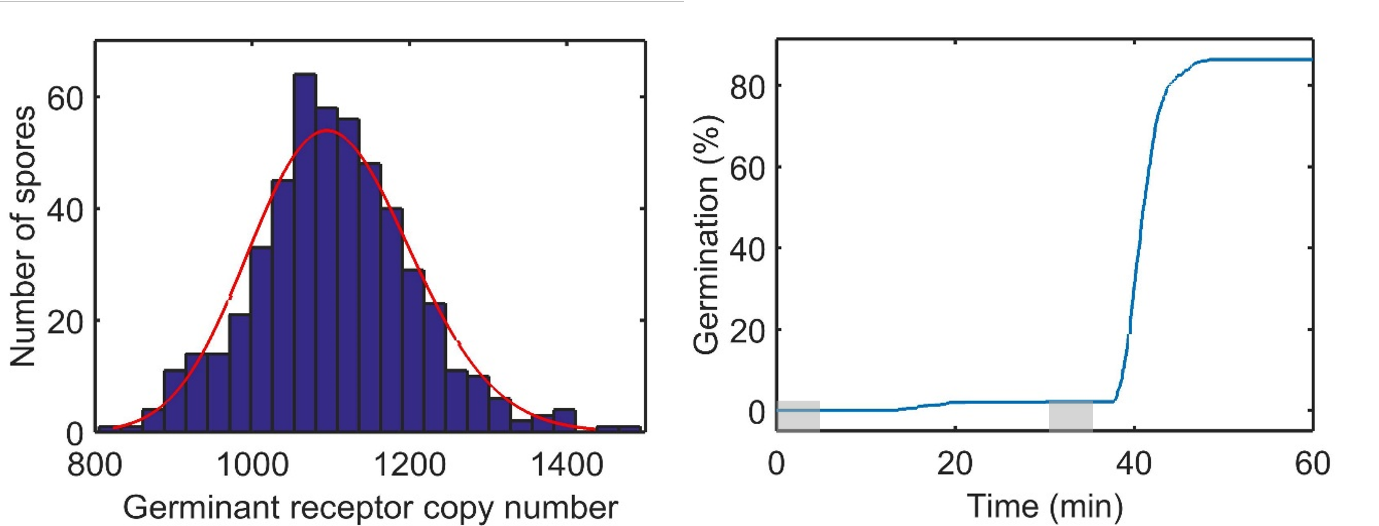

Fig S3b.	Left panel: Gamma distribution of GR copy numbers used for simulation of the percentage germination efficiency curve (Right panel). *Var*^½^(*R*_i_) is 105 at *t* is 0 min. The red line is the fitted distribution. The GR estimated mean is 1105 copies/spore, and the GR estimated standard deviation is 102 copies/spore; the parameters of the fitted gamma distribution are: *a* is 116.917 [103.304, 132.323] and *b* = 9.44706 [8.34495, 10.6947]. Right panel: Percentage germination of spores in a two pulse germination experiment.

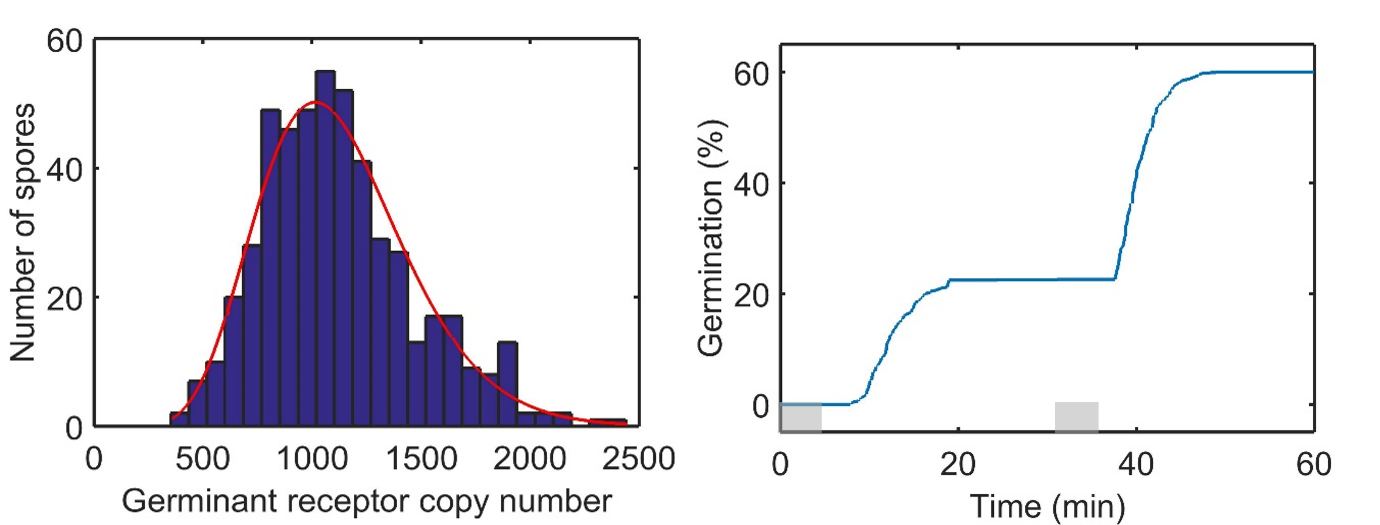

Fig S3c.	Left panel: Gamma distribution of GR copy numbers used for simulation of the germination efficiency curve (Right panel). *Var*^½^(*R*_i_) is 348 at *t* is 0 min. The red line is the fitted distribution. The GR estimated mean is 1122 copies/spore, and the GR estimated standard deviation is 346 copies/spore; the parameters of the fitted gamma distribution are: *a* is 10.4978 [9.29177, 11.8604] and *b* is 106.831 [94.2775, 121.055]. Right panel: Percentage germination of spores in a two pulse germination experiment.

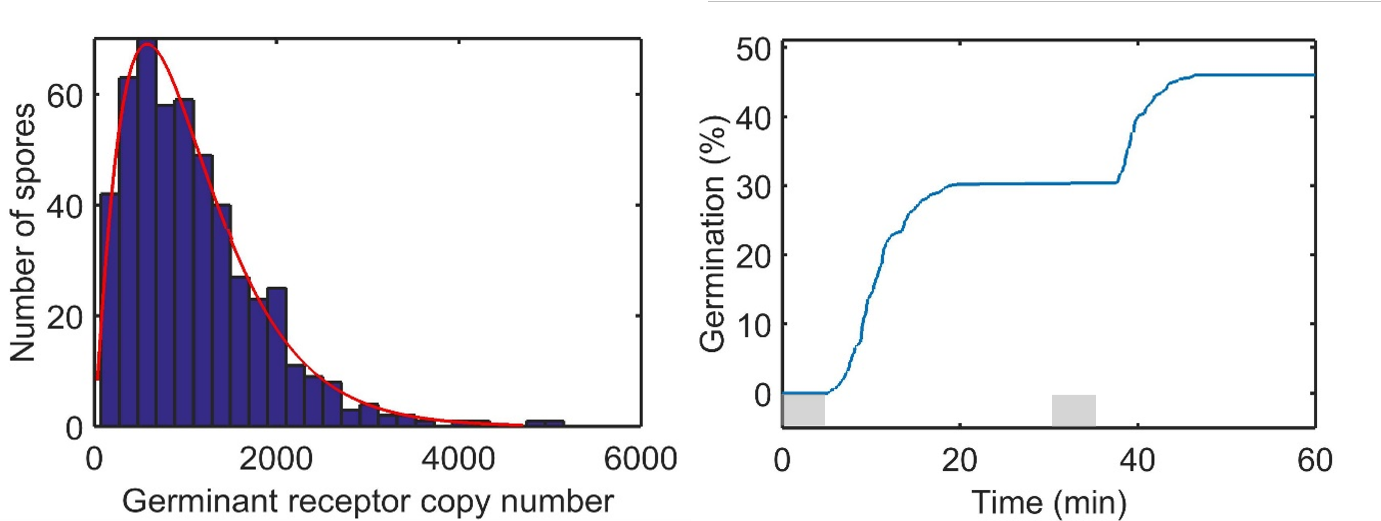

Fig S3d.	Left panel: Gamma distribution of GR copy numbers used for simulation of the percentage germination efficiency curve (Right panel). *Var*^½^(*R*_i_) is 749 at t is 0 min. The red line is the fitted distribution. The GR estimated mean is 1094 copies/spore, and the GR estimated standard deviation is 776 copies/spore; the parameters of the fitted gamma distribution are: *a* is 2.12876 [1.89643, 2.38955] and *b* is 513.649 [450.921, 585.103]. Right panel: Percentage germination of spores in a two pulse germination experiment.

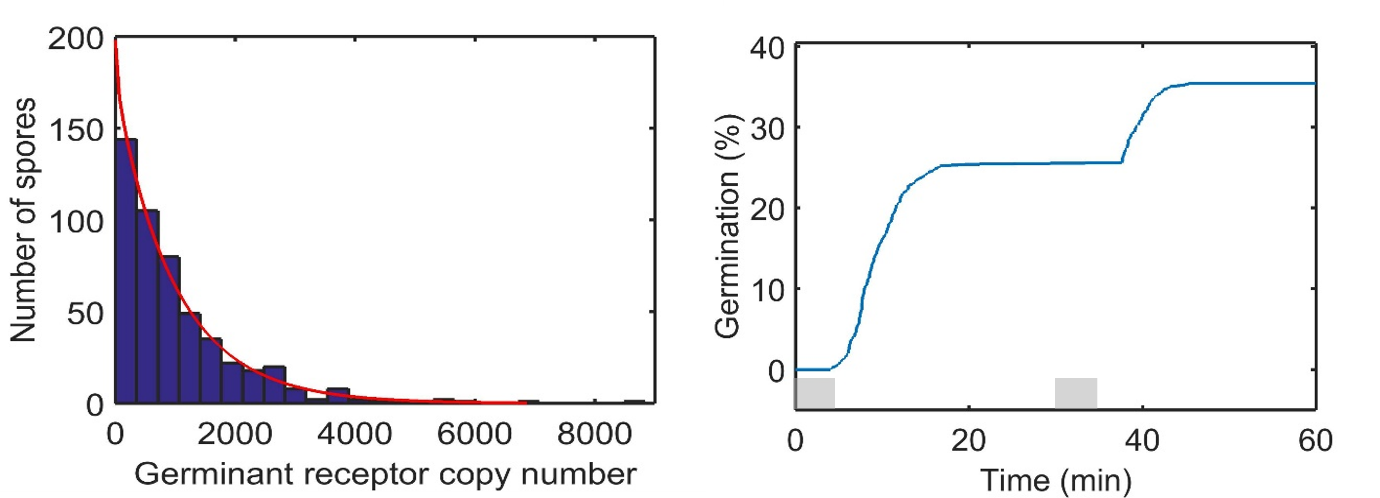

Fig S3e.	Left Panel: Gamma distribution of GR copy numbers used for simulation of the percentage germination efficiency curve. *Var*^½^(*R*_i_) is 1100 at t is 0 min. The red line is the fitted distribution. The estimated GR mean is 1023 copies/spore, and the estimated GR standard deviation is 1036 copies/spore; the parameters of the fitted gamma distribution are: *a* is 0.973473 [0.873043, 1.08546] and *b* is 1050.47 [912.749, 1208.96]. Right panel: Percentage germination of spores in a two pulse germination experiment.
