## Supplementary Figure S2 for "A mathematical kinetic model of memory in *Bacillus subtilis* spore germination"

Supplementary Material 2

The dependence of percent germination of spores on germinant concentration *S*. Germinant *S* is varied from 2.0 mM to 5.0 mM with increments of 0.5 mM. The parameters and initial values of remaining variables at time zero are *R*_a_ = 0 copies/spore; *C*_c_ = 6500 copies/spore. *C*_a_ = *C*_o_ = 0 copies/spore; *k*_1_ = 14e-3; *k*_-1_ = 100e-3; *k_2_* = 25e-7; *k*_3_ = 60e-3; *k*_4_ = 195e2; *k*_5_ = 305e-2; *n* = 3; *θ* = 20. The germinant receptor protein copy numbers are randomly drawn from a Gamma distribution with *a* = 25, *b* = 44 with *E*(*R*_i_) = 1100 and *Var*^½^(*R*_i_) = 220 at *t* is 0 min.

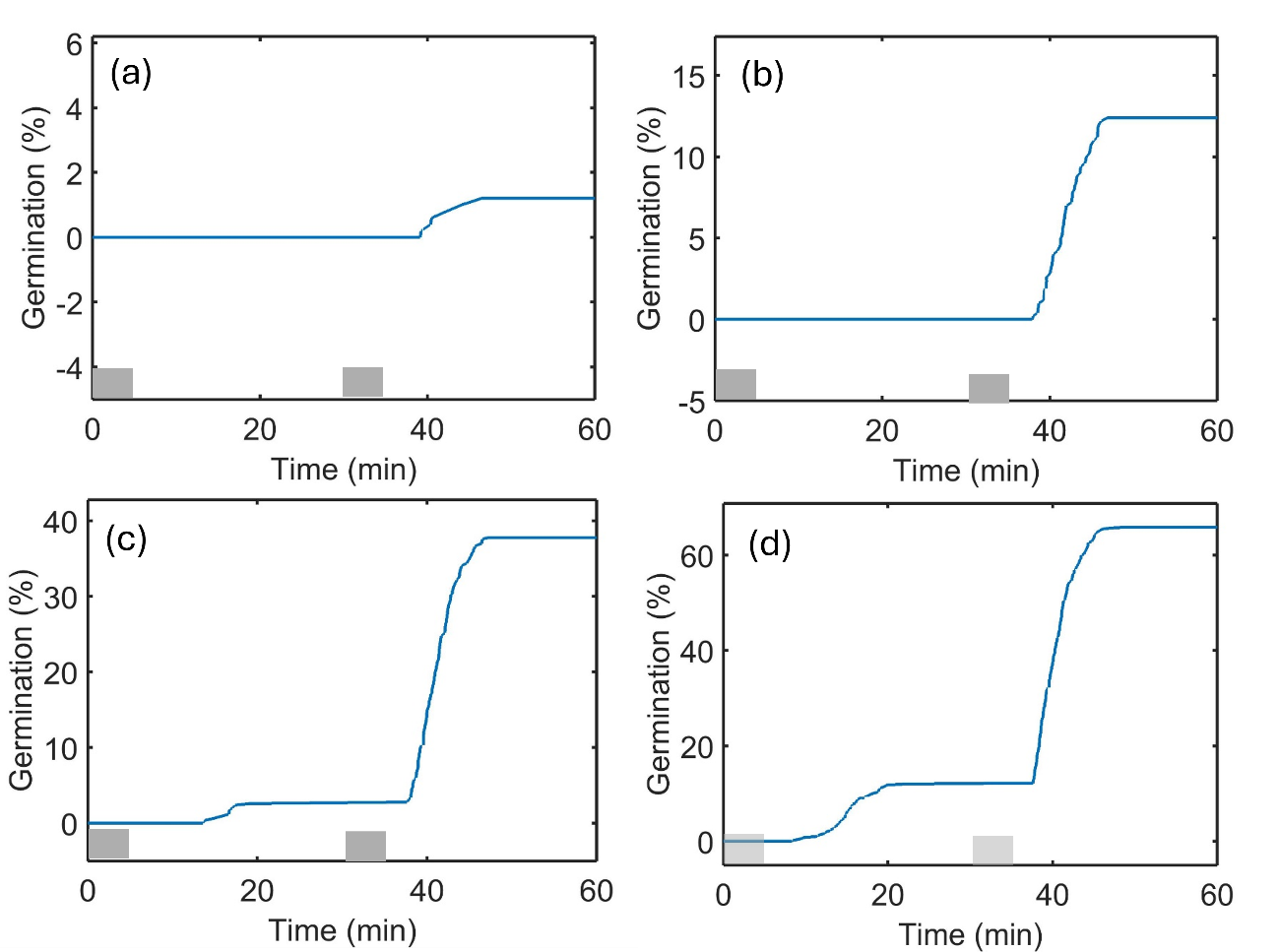

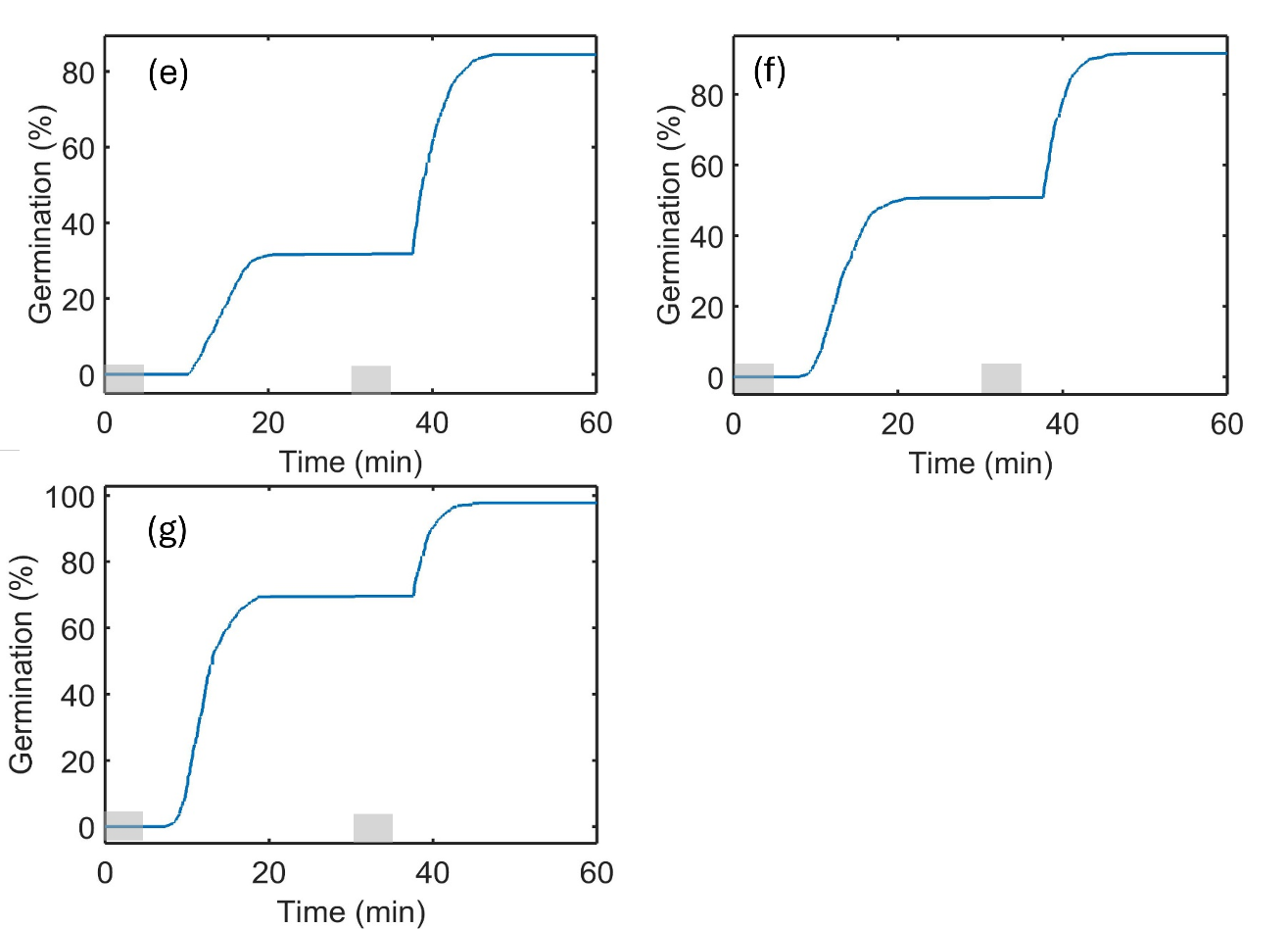

Fig.S2a-d	Percentage germination of spores in a two pulse germination experiment at (a) 2.0 mM, (b) 2.5 mM, (c) 3.0 mM, (d) 3.5 mM, (e) 4.0, (f) 4.5 mM, and (g) 5.0 mM germinant.
