## Supplementary Figure S1 for "A mathematical kinetic model of memory in *Bacillus subtilis* spore germination"

Supplementary Material 1

The effect of the GR gamma distribution mean *E*(*R*_i_) at *t* is 0 min on percentage germination in the 1^st^ and 2^nd^ germinant pulses. The scale and rate parameters *a* and *b* are varied where the standard deviation *Var*^½^(*R*_i_) at *t* is 0 min is 220 and the mean GR copy number *E*(*R*_i_) is (a) 900, (b) 1100 or (c) 1300. Two pulses of 3.5 mM germinant for receptor *R*_i_ activation are administered from 0 < *t* < 5 and 30 < *t* < 35 min. At zero time *R*_a_ = 0 copies/spore; *C*_c_ = 6500 copies/spore. *C*_a_ = *C*_o_ = 0 copies/spore; *k*_1_ = 14e-3; *k*_-1_ = 100e-3; *k_2_* = 25e-7; *k*_3_ = 60e-3; *k*_4_ = 195e2; *k*_5_ = 305e-2; *n* = 3; *θ* = 20.

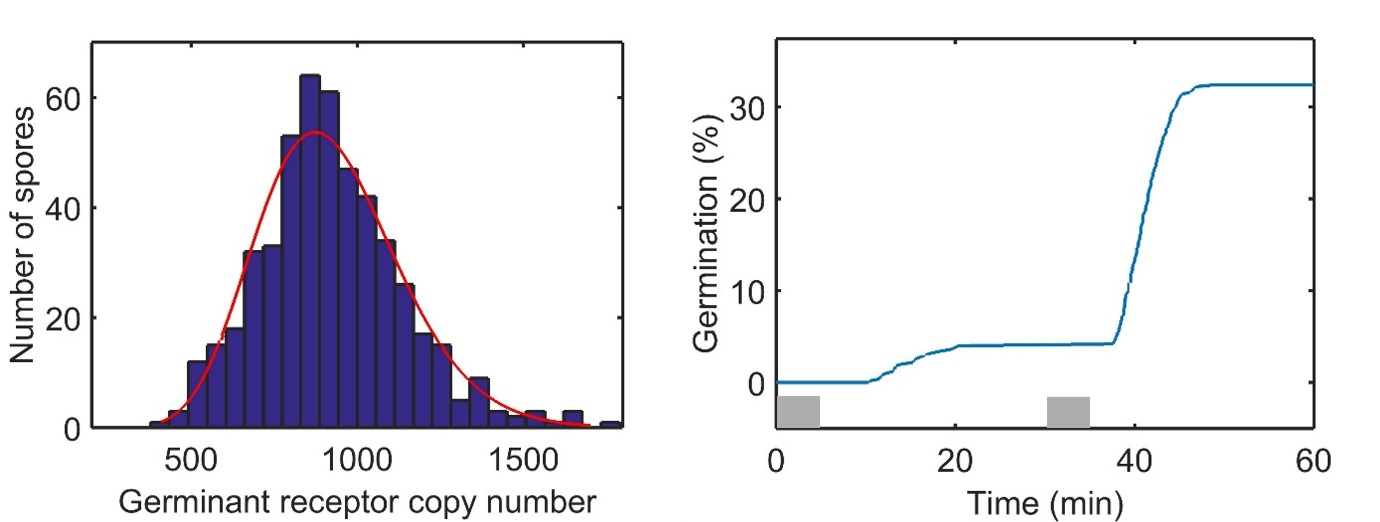

Fig S1a.	Left Panel: Gamma distribution of GR copy numbers (Left panel) used for simulation of the percentage germination efficiency curve (Right panel) with *E*(*R*_i_) is 900 at *t* is 0 min. The red line is the fitted distribution. The GR estimated mean is 924 molecules/spore, and the estimated standard deviation is 215 molecules/spore; the parameters of the fitted gamma distribution are: *a* is 18.489 [16.3515, 20.9059] and *b* is 49.9745 [44.1228, 56.6024]; Right panel: Percentage germination of spores in a two pulse germination experiment.

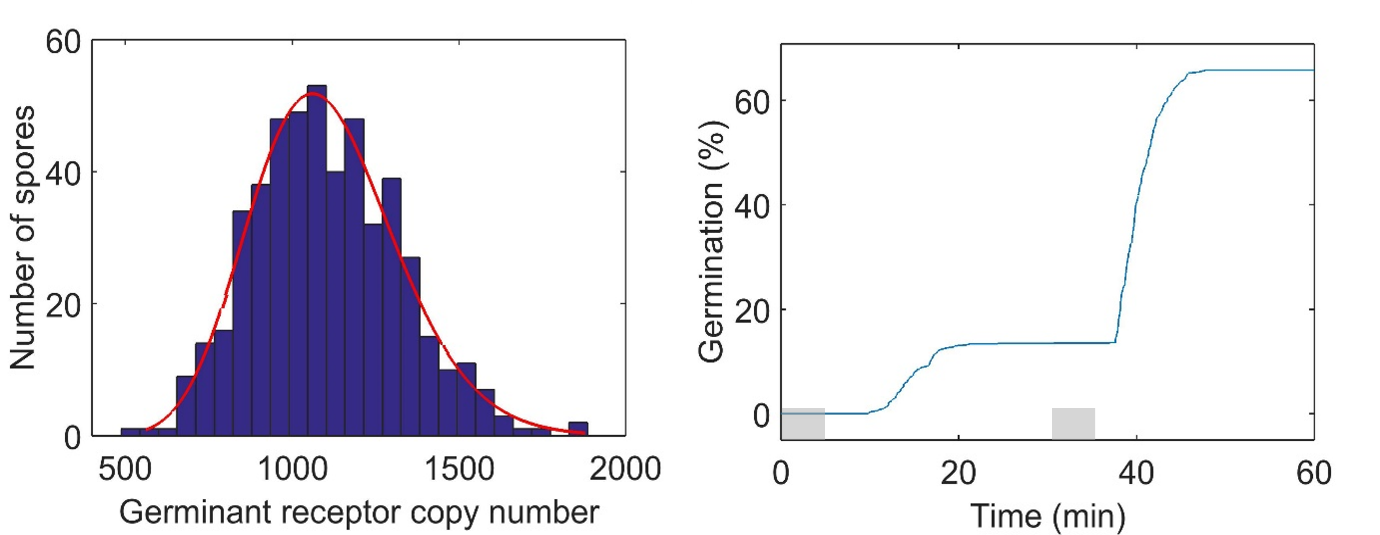

Fig S1b.	Gamma distribution of GR copy numbers (Left panel) used for simulation of the percentage germination efficiency curve (Right panel) with *E*(*R*_i_) = 1100 at *t* is 0 min. The red line is the fitted distribution. The GR estimated mean is 1105 molecules/spore, and the estimated standard deviation is 219 molecules/spore; the parameters of the fitted gamma distribution are: *a* is 25.4491 [22.5002, 28.7844] and *b* is 43.4004 [38.3247, 49.1484]; Right panel: Percentage germination of spores in a two pulse germination experiment.

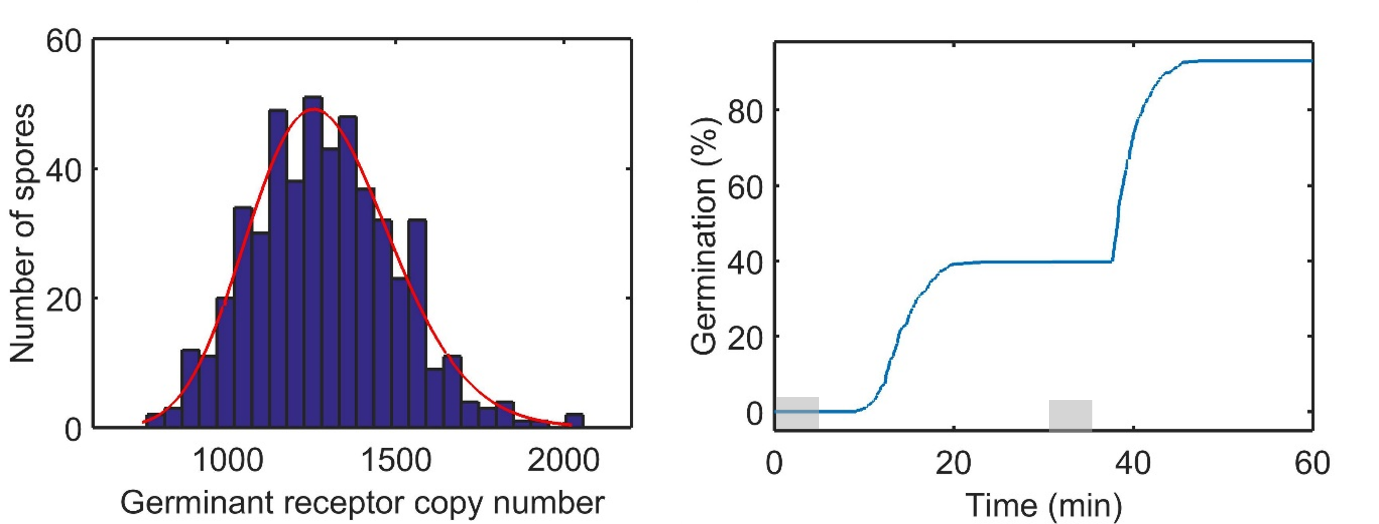

Fig S1c.	Left panel: Gamma distribution of GR copy numbers (Left panel) used for simulation of the percentage germination efficiency curve (Right panel) with *E*(*R*_i_) = 1300 at *t* is 0 min. The red line is the fitted distribution. The GR estimated mean is 1292 molecules/spore, GR estimated standard deviation is 213 molecules/spore; the parameters of the fitted gamma distribution are: *a* is 36.8424 [32.5653, 41.6813] and *b* is 35.0556 [30.9598, 39.6931]; Right panel: Percentage germination of spores in a two pulse germination experiment.
